# Stomatal CO_2_-responses at sub- and above-ambient CO_2_ levels employ different pathways in Arabidopsis

**DOI:** 10.1101/2021.05.13.443984

**Authors:** Kaspar Koolmeister, Ebe Merilo, Hanna Hõrak, Hannes Kollist

## Abstract

Stomatal pores that control plant CO_2_ uptake and water loss affect global carbon and water cycles. In the era of increasing atmospheric CO_2_ levels and vapor pressure deficit (VPD), it is essential to understand how these stimuli affect stomatal behavior. It is unknown whether stomatal responses to sub-ambient and above-ambient CO_2_ levels are governed by the same regulators and whether these responses depend on VPD. We studied stomatal conductance responses in Arabidopsis stomatal signaling mutants under conditions where CO_2_ levels were either increased from sub-ambient to ambient (400 ppm) or from ambient to above-ambient levels under normal or elevated VPD. We found that guard cell signaling components involved in CO_2_-induced stomatal closure have different roles in the sub-ambient and above-ambient CO_2_ levels. The CO_2_-specific regulators prominently affected sub-ambient CO_2_ responses, whereas the lack of guard cell slow-type anion channel SLAC1 more strongly affected the speed of above-ambient CO_2_-induced stomatal closure. Elevated VPD caused lower stomatal conductance in all and faster CO_2_-responsiveness in some studied genotypes and CO_2_-transitions. Our results highlight the importance of experimental set-ups in interpreting stomatal CO_2_- responsiveness, as stomatal movements under different CO_2_ concentration ranges are controlled by distinct mechanisms. Sometimes elevated CO_2_ and VPD responses also interact. Hence, multi-factor treatments are needed to understand plant behavior under future climate conditions.

## Introduction

Atmospheric CO_2_ concentration has nearly doubled within the past 150 years. As a result, the global average temperature has increased by 1.5°C and relative air humidity decreased across many vegetated areas (Yuan et al. 2019). Increased air temperature and decreased air humidity lead to rising values of vapor pressure deficit (VPD), the difference between actual and saturated air vapor pressures, and this increases transpiration from plants. In order to survive such conditions, plants need to adjust their water management by regulating stomatal conductance. Each stoma is composed of two guard cells and an opening formed between them; the size of the opening is controlled by increasing or decreasing guard cell turgor pressure. Guard cells adjust their turgor pressure in response to various abiotic stimuli to balance water loss and CO_2_ uptake for photosynthesis. Understanding how stomata respond to changing CO_2_ concentrations and increasing VPD is needed for breeding climate-ready crops.

Plant stomata close in response to elevated CO_2_ concentration and open in response to decreased CO_2_ concentration. Ultimately, elevated CO_2_ activates S-type anion channel SLAC1, which causes rapid stomatal closure (Vahisalu et al., 2008; Negi et al., 2008). CO_2_ can enter guard cells through the PIP2 plasma membrane channel or by transmembrane diffusion (Katsuhara and Hanba, 2008; Wang et al., 2016). Carbonic anhydrases CA1 and CA4 accelerate the conversion of intracellular CO_2_ to bicarbonate (HCO_3_^-^), which can act as a second messenger (Hu et al., 2010). In guard cells, CO_2_/HCO_3_^-^ promotes interaction between the protein kinases MPK4/MPK12 and the Raf-like kinase HT1, leading to HT1 inhibition, which is an essential step in the regulation of stomatal responses to CO_2_ (Hashimoto et al., 2006; Hashimoto-Sugimoto et al., 2016; Hõrak et al., 2016; Jakobson et al., 2016; Takahashi et al., 2022; Yeh et al., 2023). HT1 phosphorylates the CBC1/CBC2 Raf-like kinases that function downstream of HT1 and these Raf kinases can inhibit the S-type anion channel activation via a currently unknown mechanism (Hõrak et al., 2016; Hiyama et al., 2017; Hayashi et al., 2020). Stomata in HT1-deficient plants do not respond to CO_2_ concentration changes while carbonic anhydrase, MPK12 and SLAC1 mutants exhibit impaired stomatal CO_2_ responses (Hashimoto et al., 2006; Vahisalu et al., 2008; Hu et al., 2010; Hashimoto-Sugimoto et al., 2016; Hõrak et al., 2016; Jakobson et al., 2016).

Elevated VPD increases transpiration and reduces epidermal turgor that due to mechanical interactions between guard and epidermal cells in angiosperms leads to faster light-induced stomatal opening (Mott et al., 1999; Pichaco et al., 2024). To prevent wilting, stomata close under elevated VPD. Abscisic acid (ABA) is a drought-induced plant stress hormone and an important stomatal regulator. VPD has a direct effect on ABA concentration as increased ABA levels in angiosperms were observed 20 minutes after increasing VPD from 0.7 to 1.5 kPa (McAdam and Brodribb, 2015). Considering that ABA has a major role in angiosperm stomatal closure (Cutler et al., 2010), elevated VPD might prime plant stomata for fast closure via increased ABA concentration. The protein kinase OST1 and the leucine-rich receptor-like pseudokinase GHR1 are activated in the presence of ABA and trigger anion efflux through the major guard cell slow-type anion channel SLAC1 (Brandt et al., 2012; Hua et al., 2012; Sierla et al., 2018). OST1 and GHR1 are both involved in elevated VPD- and CO_2_-induced stomatal closure response (Xue et al., 2011; Merilo et al., 2018; Sierla et al., 2018; Hsu et al., 2021; Jalakas et al., 2021b).

Understanding CO_2_-induced plant stomatal closure responses is essential for future plant breeding. Due to changing climate conditions, it is also important to understand if and how stomatal CO_2_ regulation is affected by elevated VPD levels. Previous work in grasses suggests that elevated VPD levels reduce both stomatal conductance and stomatal sensitivity to CO_2_ concentration changes (Morison and Gifford, 1983) but the interactions of CO_2_ and humidity responses in dicots remain poorly understood.

Under light, CO_2_ concentration inside the leaf is usually below ambient levels and this causes stomatal opening. Stomata close when CO_2_ concentration inside the leaf increases to ambient levels, and an additional rise in CO_2_ concentration to above-ambient levels causes further stomatal closure (Brodribb et al., 2009; Hõrak et al., 2017). Thus, stomatal closure response exists within sub-ambient as well as in above-ambient CO_2_ levels, however it is not clear whether these responses are controlled by the same regulators. In some studies, CO_2_-induced stomatal closure is defined as the response to an increase in CO_2_ concentration from ambient to above-ambient levels (Franks and Britton-Harper, 2016; Hõrak et al., 2017), while some studies use a [CO_2_] change from sub-ambient to above-ambient levels (Azoulay-Shemer et al., 2015; Chater et al., 2015). Data from previous studies comparing CO_2_ responses in ferns and angiosperms suggests that stomatal responses to CO_2_ are different, when changing CO_2_ levels in the sub-ambient or above-ambient ranges (Brodribb et al., 2009; Hõrak et al., 2017). To address the underlying mechanisms of CO_2_-induced stomatal closure at different CO_2_ transitions under normal and elevated VPD conditions, we studied plants deficient either in guard cell anion channel SLAC1 and its activation (*slac1-3*, *ghr1-3*, *ost1-3*) or in the CO_2_-specific stomatal signaling branch regulated by MPK12 and HT1 kinases (*mpk12-4*, *ht1-2*, *ht1-8D*) and carbonic anhydrases CA1 and CA4 (*ca1ca4*). Our results show that different stomatal regulators have a different degree of importance in CO_2_-induced stomatal closure in sub-ambient and above-ambient CO_2_ levels and are also differently affected by VPD.

## Materials and Methods

### Plant lines and growth conditions

*Arabidopsis thaliana* accession Col-0 and the following mutants in the same genetic background were used for experiments: *slac1-3* (Vahisalu et al., 2008), *ost1-3* (Yoshida et al., 2002), *ghr1-3* (Sierla et al., 2018), *ht1-2* (Hashimoto et al., 2006), *ht1-8D* (Hõrak et al., 2016), *mpk12-4* (Jakobson et al., 2016), *ca1ca4* (Hu et al., 2010). Plants were grown in 4:2:3 v/v peat:vermiculite:water mixture at 12/12 photoperiod with 150 µmol m^-2^ s^-1^ light in controlled-environment growth cabinets (AR-66LX; Percival Scientific; MCA1600, Snijders Scientific) at 70% relative humidity and day-time temperature of 23°C (VPD 0.84 kPa) and night-time temperature 18°C (VPD 0.62 kPa). Plant age at experiment time was ∼25 days.

### Gas-exchange measurements

Measurements of stomatal conductance were carried out with a temperature-controlled custom-built gas-exchange device (Kollist et al., 2007; Hõrak et al., 2017). Plants were inserted into measurement cuvettes and allowed to acclimate for 1-2 hours at normal air humidity (VPD, 0.9 kPa) or at lower air humidity (VPD, 2.3 kPa), 24°C air temperature and 400 ppm CO_2_. Experiments with various CO_2_ transitions were carried out and in some cases both under normal and at high VPD conditions, as shown in Fig. 1. The first CO_2_ treatment was applied approximately at noon (11:30 to 12:30), stomatal conductance was always followed for 2 hours under each treatment (Fig. 1).

**Figure 1.**
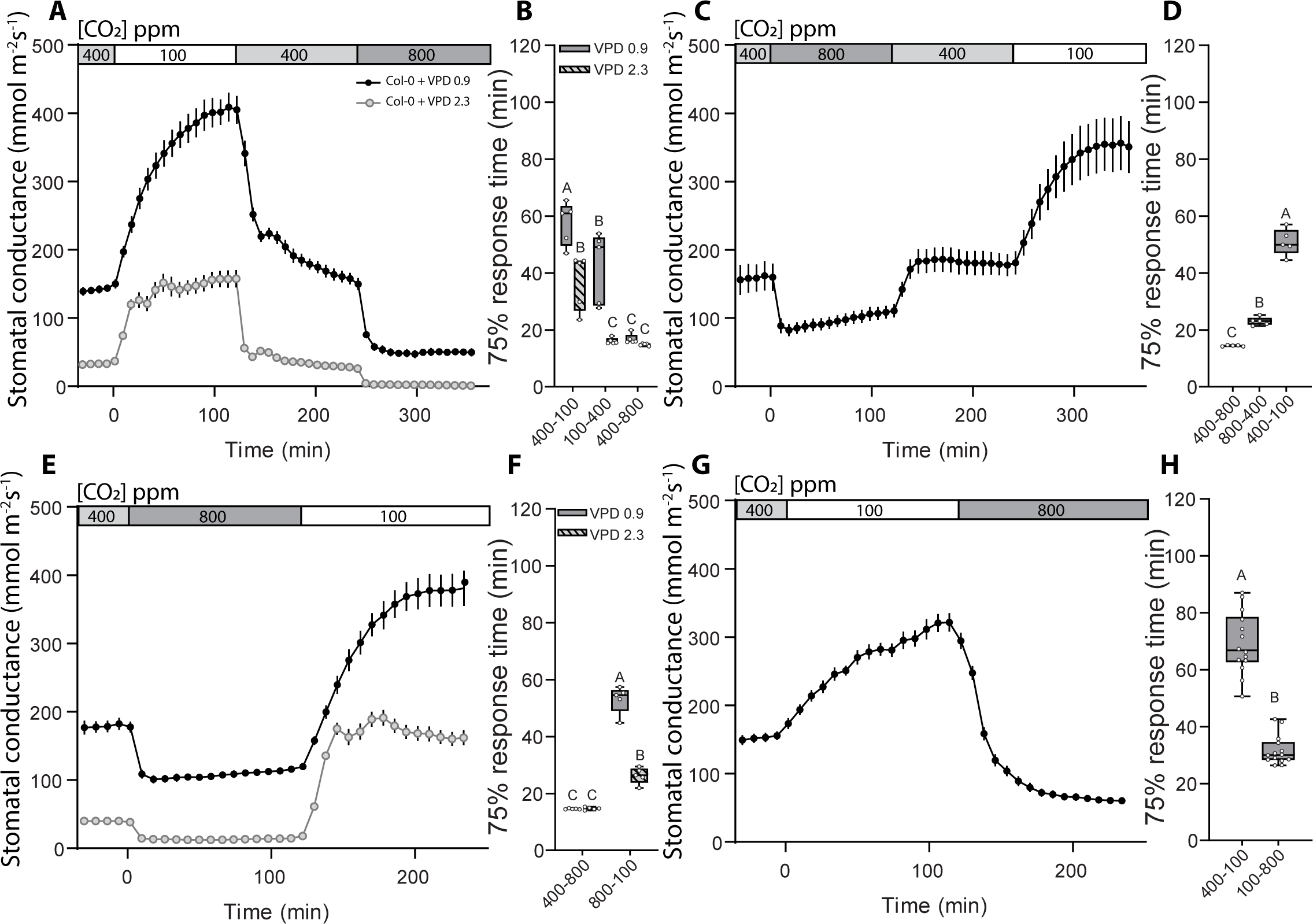
Kinetics of CO_2_-induced stomatal responses in wild-type Arabidopsis with regular and elevated VPD. Col-0 wild-type Arabidopsis stomatal response to sequential changes in CO_2_ concentration under regular **(A,C,E,G)** and elevated **(A,E)** VPD conditions. Mean stomatal conductance ± SEM is shown. **(B,D,F,H)** Boxplot of 75% response time (minutes) of stomatal response to CO_2_ concentration changes. Boxes represent 25-75 % quartiles and median as the horizontal lines, whiskers indicate the smallest and largest values, points show individual plant values. **(B,F)** Statistically significantly different groups are indicated with different letters (Two-way ANOVA with Tukey *post-hoc* test). **(D,H)** Statistically significantly different groups are indicated with different letters (One-way ANOVA with Tukey *post-hoc* test). Sample size was 5 in **(A,B,C,D,E,F)** and 14 in **(G,H)**. Start of first treatment was between 11:30 to 12:30.

### Data analysis

Magnitude of stomatal closure response was calculated as the absolute difference in stomatal conductance between last time point before treatment and at the end of 2 hours of treatment at a given CO_2_ level (displayed as ‘Change in g_s_^’^). To describe the kinetic characteristics of stomatal responses, we calculated the 75% response time by scaling the two-hour stomatal responses to a range from 0% to 100% and by calculating the time when 75% of the total stomatal response was achieved. One-way ANOVA with Tukey *post hoc* test was used for statistical analysis as indicated in the figure legends, p < 0.05 were considered statistically significant. Statistical analyses were carried out with Past 4.0 (Hammer et al., 2001) and Statistica 7.1 (Stat. Soft. Inc).

## Results

### Stomatal closure kinetics are different between sub-ambient to ambient and ambient to above-ambient [CO_2_] transitions

We analyzed stomatal responses to CO_2_ in the sub-ambient and above-ambient concentration ranges in the model plant *Arabidopsis thaliana* to clarify whether these responses are controlled by the same or by different regulators. Four different CO_2_ transition sequences were used (Fig. 1A,C,E,G); in two set-ups we applied high VPD (2.3 kPa) as an additional factor before and throughout CO_2_ treatments (Fig. 1A,E). Experiments were started with ambient (400ppm) CO_2_ concentration and each consecutive CO_2_ treatment lasted for 2 hours. This approach allowed to investigate plant stomatal response to different CO_2_ transitions – stomatal opening in response to CO_2_ transition from 400 to 100 ppm (from here on referred to as 400-100) (Fig. 1A,C,G), from 800 to 400 ppm (800-400) (Fig. 1C), or from 800 to 100 ppm (800-100) (Fig. 1E), and stomatal closure in response to CO_2_ transition from 100 to 400 ppm (100-400) (Fig. 1A), from 400 to 800 ppm (400-800) (Figure 1A,C,E), or from 100 to 800 ppm (100-800) (Fig. 1G).

Stomatal closure responses had clearly different 75% response time in Col-0 wild-type plants at different CO_2_ transitions: the 400-800 stomatal closure was faster than 100-400 closure under normal VPD (Fig. 1A,B). The rapid 400-800 response was consistent throughout all experimental setups (Fig. 1A-F). The observed different kinetics of these CO_2_-induced stomatal closure responses suggest that they could be regulated by different components. Stomatal opening responses to sub-ambient CO_2_ levels, 400-100 (Fig. 1A-D,G,H) and 800-100 (Fig. 1E,F) had relatively slow 75% response times, whereas the 800-400 response, opening from above-ambient to ambient CO_2_ levels, had lower response range and faster 75% response time (Fig. 1C,D). This suggests that stomatal opening to sub-ambient CO_2_ concentrations is a much stronger, albeit slower, response than stomatal opening during the above-ambient to ambient [CO_2_] change.

### High VPD but not the order of CO_2_ transitions affects stomatal CO_2_ response kinetics

To study whether high VPD affects CO_2_ responses, we conducted the 400-100-400-800 and the 400-800-100 CO_2_ transitions under conditions where plants were first acclimatized to increased VPD (2.3 kPa) for approximately three hours and then subjected to changes in CO_2_ levels under the elevated VPD conditions. Under such conditions, both the 400-100 and 800-100 stomatal opening responses were significantly faster than under normal VPD (Fig. 1A,B E,F). Stomatal closure in sub-ambient to ambient CO_2_ concentration range was also enhanced under high VPD conditions, resulting in shorter stomatal 75% response time during the 100-400 ppm [CO_2_] transition compared with normal VPD (Fig. 1A,B), whereas 400-800 ppm [CO_2_] transition 75% response time was not affected by high VPD (Fig. 1A,B,E,F). These results suggest that during high VPD stress, plant stomata could be primed for faster movements in the sub-ambient to ambient CO_2_ concentration range, but VPD has a minor role in above-ambient CO_2_-induced stomatal closure.

In our first experiments, the 400-100 opening stimulus was applied before the 400-800 closure. To test whether the exposure to sub-ambient CO_2_ levels had an effect on stomatal CO_2_-responsiveness, we also applied the CO_2_ transitions in reverse order (400-800-400-100, Fig. 1C). Stomatal 75% response times during the 400-100 transition appeared slightly shorter when it was the last transition (Fig. 1C,D) than when it was the first (Fig. 1A,B,G,H), whereas the 400-800 transition response speed was unaffected by previous treatments (Fig. 1A,B,C-F). Thus, there were no major effects of the order of CO_2_ concentration transitions on stomatal CO_2_-response kinetics.

### The CO_2_-specific pathway components MPK12 and carbonic anhydrases are more involved in sub-ambient to ambient than in above-ambient CO_2_-induced stomatal closure

To better understand the role of the CO_2_-signalling module comprising MPK12, HT1 and carbonic anhydrases CA1 and CA4 in stomatal CO_2_ responses at different CO_2_ levels, we analyzed the *mpk12-4*, *ht1-2*, *ht1-8D*, and *ca1a4* in a similar experimental set-up as described for wild-type plants in Fig. 1A. All the CO_2_-signalling mutants showed very little stomatal closure in magnitude compared with wild-type plants, whereas 75% response time was significantly reduced only in the *ca1ca4* mutant in response to the 100-400 transition (Fig. 2A,C,E). The 75% response times for the *ht1-2* and *ht1-8D* mutants in this CO_2_ transition are not informative due to hardly any stomatal response in these mutants (Fig 2A,C,E). The response of *mpk12-4* to the 100-400 transition was as fast as in wild-type, but significantly smaller in magnitude (Fig. 2A,C,E). In response to the 400-800 transition, the HT1 mutants had very weak stomatal response with small magnitude and slow speed, whereas stomatal closure in the *mpk12-4* and *ca1ca4* plants was slower than in WT, but larger in magnitude (Fig. 2A,C,E). Very weak responses of the HT1 mutants suggest that HT1 is necessary to initiate stomatal responses to CO_2_ concentration changes in both ambient and sub-ambient levels. MPK12 and carbonic anhydrase mutants display stronger stomatal responses in the above-ambient than sub-ambient CO_2_ range (Fig. 2A,C,E), suggesting that respective signaling components have a more important role at sub-ambient compared with above-ambient CO_2_ concentrations.

**Figure 2.**
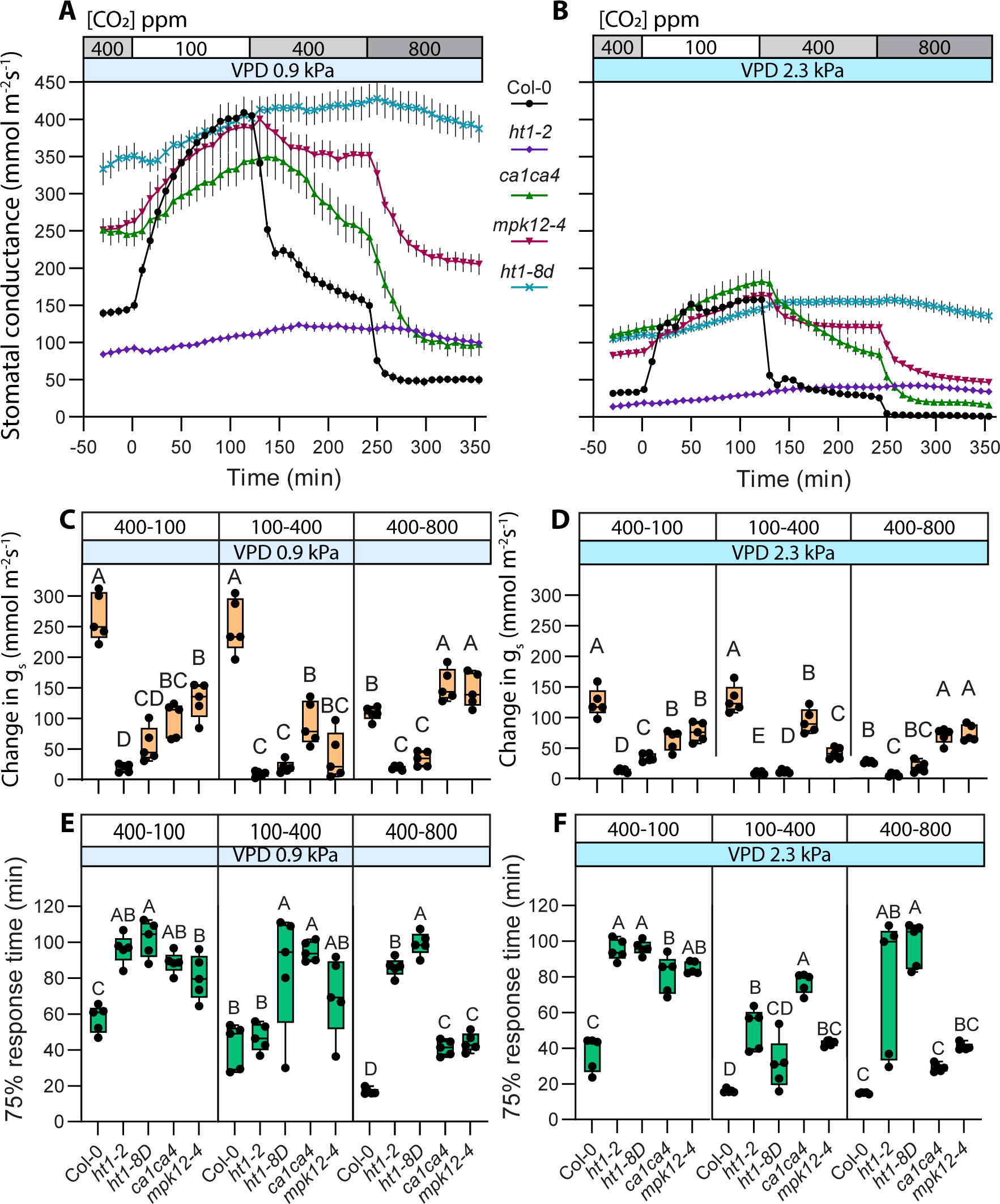
CO_2_ pathway mutants retain response patterns under regular and elevated VPD conditions. **(A)** and **(B)** Stomatal response to CO_2_ concentration changes from 400 to 100 ppm, 100 to 400 ppm and 400 to 800 ppm under regular **(A)** and elevated **(B)** VPD conditions, mean stomatal conductance ± SEM is shown. **(C,D)** Boxplot of stomatal conductance (g_s_) change (mmol m^-2^ s^-1^) in response to CO_2_ concentration changes from 400 to 100 ppm, 100 to 400 ppm and 400 to 800 ppm, respectively. **(E,F)** Boxplot of 75% response time (minutes) of stomatal response to CO_2_ concentration changes from 400 to 100 ppm, 100 to 400 ppm and 400 to 800 ppm, respectively. **(C-F)** Boxes represent 25-75 % quartiles and median as the horizontal lines, whiskers indicate the smallest and largest values, points show individual plant values. Statistically significantly different groups are marked with different letters (One-way ANOVA with Tukey *post hoc* test). **(A-F)** Sample size was 5 for all plant lines. VPD during experiments was 0.9 kPa in **(A,C,E)** and 2.3 kPa in **(B,D,F)**. Start of first treatment was between 11:30 to 12:30. Col-0 data is same as used in figure 1, experiments with Col-0 and mutant lines were done together.

Elevated VPD led to lower stomatal conductance in all studied mutants (Fig. 2A,B), and thus smaller magnitudes of CO_2_-induced changes in stomatal conductance (Fig. 2C,D). Elevated VPD had some effects on the patterns of CO_2_-responsiveness in the CO_2_-signalling mutants (Fig 2A-F). The *ca1ca4* and *mpk12-4* stomatal response times were similar to WT in the 400-800 transition under elevated VPD (Fig. 2F), and the *mpk12-4* mutant had a significantly slower than wild-type response to the 100-400 transition only under elevated VPD. Thus, the relatively larger degree of importance of MPK12 and carbonic anhydrases in regulating stomatal responses in the sub-ambient CO_2_ ranges was more pronounced under elevated VPD conditions.

Stomatal opening response to 400-100 transition in all of the studied CO_2_-signalling mutants was lower in magnitude and slower in response time, irrespective of VPD, whereas the difference in response time between wild-type and mutants was larger under elevated VPD (Fig. 2A-F). Thus, stomatal opening response to sub-ambient [CO_2_] is affected by all of the CO_2_ signaling pathway components represented in this study.

### SLAC1 and GHR1 are more important for above-ambient CO_2_-induced stomatal closure

The SLAC1 anion channel is a major component in the activation of stomatal closure response. Thus, we examined CO_2_ responses across different CO_2_ concentration ranges in plants deficient in the SLAC1 anion channel (*slac1-3*) and in OST1 or GHR1: proteins involved in SLAC1 activation (*ost1-3*, *ghr1-3*, Fig. 3). Stomatal 75% response time of *slac1-3* plants was longer in both, 100-400 and 400-800 CO_2_ transitions (Fig. 3A,E) but the latter was more affected as the 75% response time difference compared with wild-type plants was notably larger in the 400-800 transition. Yet, the magnitude of stomatal closure between wild-type Col-0 and *slac1-3* was only different in the 100-400 transition (Fig. 3C), while their 400-800 response magnitude was similar. Under elevated VPD, *slac1-3* stomatal response amplitude was similar to wild-type both during the 100-400 and 400-800 transitions, although stomatal response times of *slac1-3* were still significantly longer than wild-type on both transitions (Fig. 2D,F). Together, these results show that SLAC1 is important in stomatal closure in both sub-ambient and above-ambient [CO_2_] ranges, but response speed tends to be more severely impacted in the 400-800 transition, suggesting a more prominent role for SLAC1 in ensuring fast above-ambient CO_2_-induced stomatal closure.

**Figure 3.**
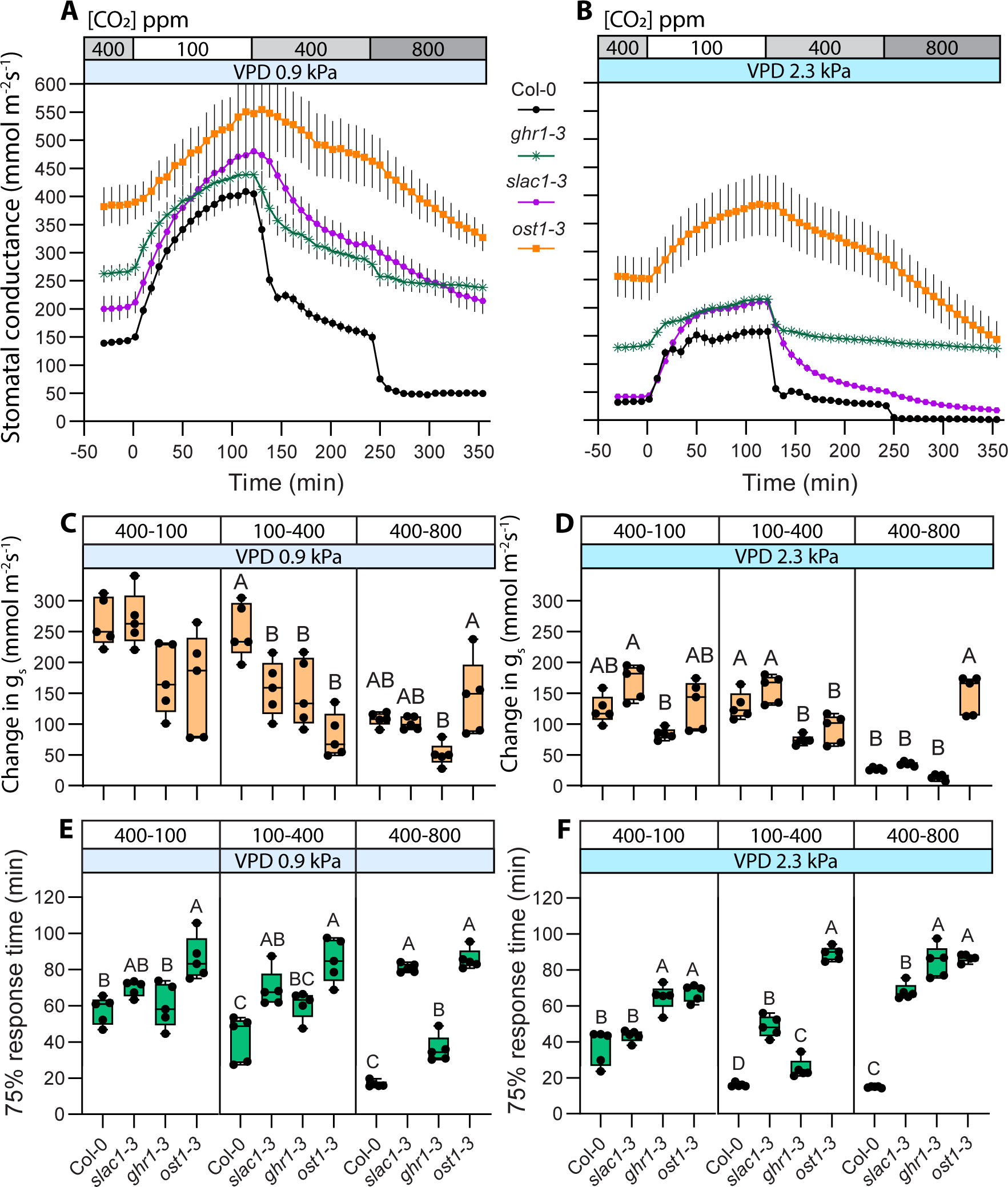
CO_2_ response patterns in anion channel activation mutants are affected by VPD conditions. **(A)** and **(B)** Stomatal response to CO_2_ concentration changes from 400 to 100 ppm, 100 to 400 ppm and 400 to 800 ppm in regular **(A)** and elevated **(B)** VPD conditions, mean stomatal conductance ± SEM is shown. **(C,D)** Boxplot of stomatal conductance (g_s_) change (mmol m^-2^ s^-1^) in response to CO_2_ concentration changes from 400 to 100 ppm, 100 to 400 ppm and 400 to 800 ppm, respectively. **(E,F)** Boxplot of 75% response time (minutes) of stomatal response to CO_2_ concentration changes from 400 to 100 ppm, 100 to 400 ppm and 400 to 800 ppm, respectively. **(C-F)** Boxes represent 25-75 % quartiles and median as the horizontal lines, whiskers indicate the smallest and largest values, points show individual plant values. Statistically significantly different groups are marked with different letters (One-way ANOVA with Tukey *post hoc* test). **(A-F)** Sample size was 5 for all plant lines. VPD during experiments was 0.9 kPa in **(A,C,E)** and 2.3 kPa in **(B,D,F)**. Start of first treatment was between 11:30 to 12:30. Col-0 data is same as used in figure 1, experiments with Col-0 and mutant lines were done together.

Plants deficient in SLAC1 activation via OST1 or GHR1 also showed impaired CO_2_- induced stomatal closure both in the 100-400 and 400 to 800 transitions (Fig. 3A,C,E). The *ost1-3* mutant had long stomatal response times in both the 100-400 and 400-800 transitions, whereas the *ghr1-3* response was similar to wild-type in the 100-400 transition, but slower and very small in magnitude during the 400-800 transition (Fig. 3A,C,E). Thus, both SLAC1-activating proteins are involved in CO_2_ responses in all tested concentration ranges, but GHR1, like SLAC1, appears to contribute more towards above-ambient CO_2_-induced stomatal closure.

As in other mutants (Fig. 2), elevated VPD lowered stomatal conductance and stomatal response magnitude, but tended to increase stomatal opening speed (Fig. 3). The *ost1-3* 75% response times and magnitude remained similar in both VPD conditions, in line with its VPD-insensitivity (Fig 3C-F, Merilo et al., 2013). In *ghr1-3*, stomatal response to 400-800 was either absent or extremely weak under elevated VPD (Fig. 3B,D).

Sub-ambient CO_2_-induced stomatal opening response magnitude was similar to wild-type in all of the studied anion channel activation mutants, irrespective of VPD (Fig 3C,D). Stomatal 75% response time under regular VPD was similar to wild-type in all but *ost1-3* (Fig. 3E), whereas under elevated VPD conditions both *ghr1-*3 and *ost1-*3 had longer stomatal opening 75% response times (Fig. 3F). Therefore, regulation of SLAC1 is less important for sub-ambient CO_2_-induced stomatal opening than the guard cell CO_2_-specific signaling pathway. Only under elevated VPD, the 400-100 transition response was slower compared to wild-type in *ghr1-3* (Fig. 3F), indicating an interaction between CO_2_ and VPD signaling in stomatal opening responses.

### Shifting CO_2_ levels from 100 to 800 ppm masks the differences between stomatal behavior in 100-400 and 400-800 ppm [CO_2_] transitions

To further study how stomatal movements differ depending on CO_2_ concentrations, we did additional gas exchange measurements, during which CO_2_ concentration was changed directly from sub-ambient (100 ppm) to above-ambient (800 ppm). Previously, in Fig. 2 and 3 we saw different stomatal response characteristics for 100-400 and 400-800 CO_2_ transitions for different mutants that we failed to detect during the 100-800 CO_2_ transition (Fig. 4). For example, *mpk12-4* stomatal response was small in amplitude in the 100-400 transition, but much larger in the 400-800 transition (Fig. 2A,C), whereas in the 100-800 transition the *mpk12-4* stomatal response amplitude was similar to wild-type (Fig. 4A). Similarly, in the *ca1ca4* plants 100-400 stomatal response was smaller and 400-800 response larger than in wild-type in amplitude (Fig. 2C), yet the 100-800 response magnitude was similar to wild-type (Fig. 4C). The *ghr1-3* mutant had slower response to the 400-800 transition (Fig. 3E), whereas its 75% response time was similar to wild-type in the 100-800 transition (Fig. 4F). These results demonstrate that differences in stomatal responses between plant lines can remain undiscovered depending on CO_2_ concentrations that are used for experiments.

**Figure 4.**
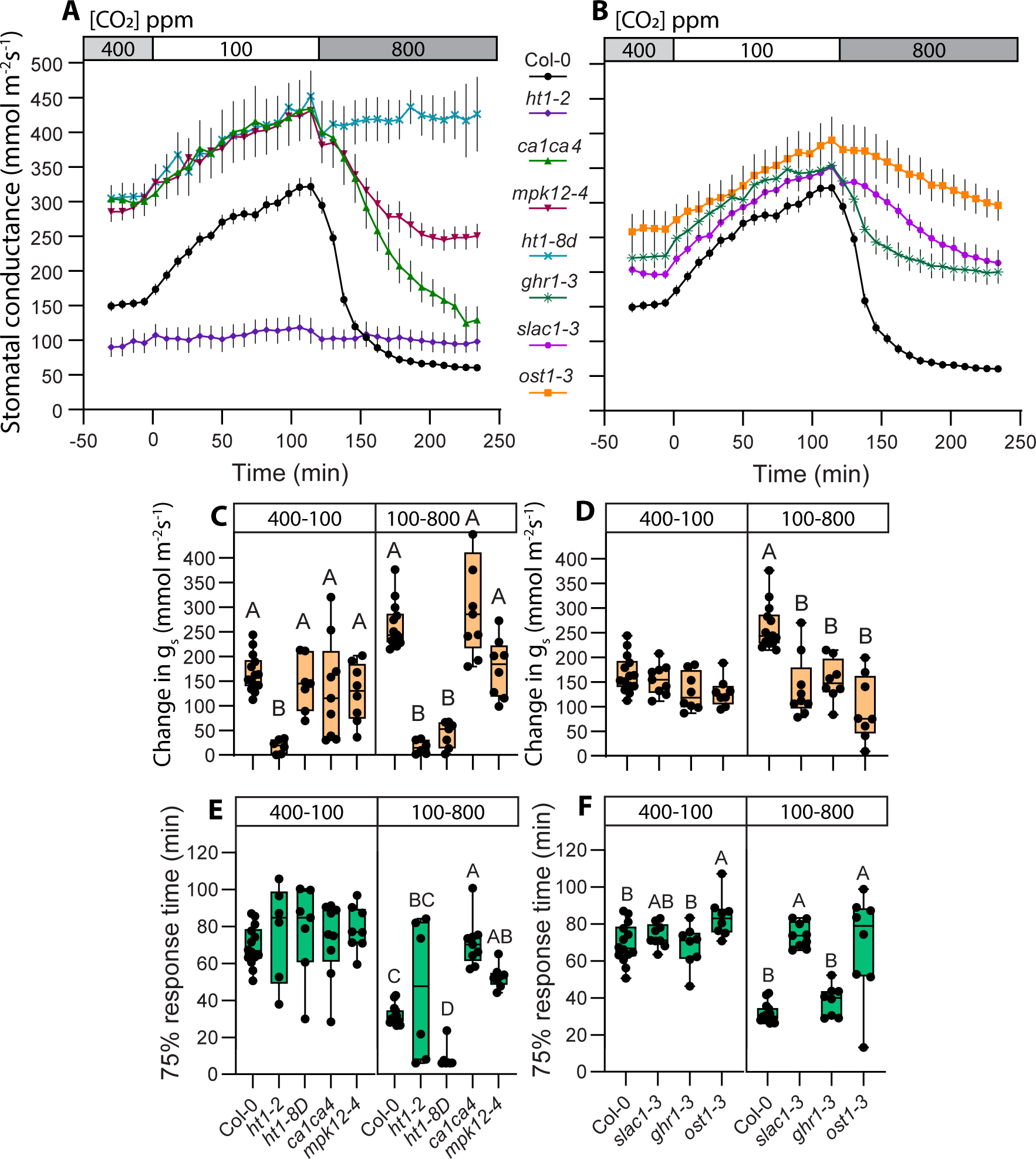
CO_2_ transition from 100 to 800 ppm masks different responses present in 100 to 400 and 400 to 800 ppm CO_2_ transitions. **(A)** and **(B)** Stomatal response to CO_2_ concentration changes from 400 to 100 ppm and 100 to 800 ppm, mean stomatal conductance ± SEM is shown. **(C,D)** Boxplot of stomatal conductance (g_s_) change (mmol m^-2^ s^-1^) in response to CO_2_ concentration changes from 400 to 100 ppm and 100 to 800 ppm, respectively. **(E,F)** Boxplot of 75% response time (minutes) of stomatal response to CO_2_ concentration changes from 400 to 100 ppm and 100 to 800 ppm, respectively. **(C-F)** Boxes represent 25-75 % quartiles and median as the horizontal lines, whiskers indicate the smallest and largest values, points show individual plant values. Statistically significantly different groups are marked with different letters (One-way ANOVA with Tukey *post hoc* test). **(A-F)** Sample size was 14 for Col-0; 6 for *ht1-2*; 7 for *ht1-8D*; 8 for *ghr1-3*, *mpk12-4* and *ost1-3*; 9 for *slac1-3* and *ca1ca4*. VPD during experiments was 0.9 kPa in **(A-F)**. Start of first treatment was between 11:30 to 12:30. Col-0 data is same as used in figure 1, experiments with Col-0 and mutant lines were done together.

Similar to CO_2_-induced stomatal closure experiments, [CO_2_] ranges were important also in stomatal opening assays (Fig. 2-3 vs Fig. 5). For example, response to the 400-100 transition in *mpk12-4* was slower and smaller in magnitude compared with wild-type plants (Fig 2C,E), but in the 800-100 experiments, *mpk12-4* had normal response amplitude and wild-type-like 75% response time (Fig. 5C,E). The *slac1-3* plants had similar to wild-type 400-100 stomatal opening speed and magnitude (Fig. 3C,E), but were slower in the 800-100 response (Fig. 5F). We also observed a significantly slower opening response to the 800-400 transition in *slac1-3* (Supplementary Fig. 1F), which indicates that the reduced stomatal opening speed of the 800-100 response in this mutant is caused by slower opening in the 800-400 range. Thus, although wild-type plants have no discernible differences between the 400-100 and the 800-100 stomatal opening responses (Fig. 1A,B,E,F), the molecular mechanisms are at least partly different for these CO_2_ transitions.

**Figure 5.**
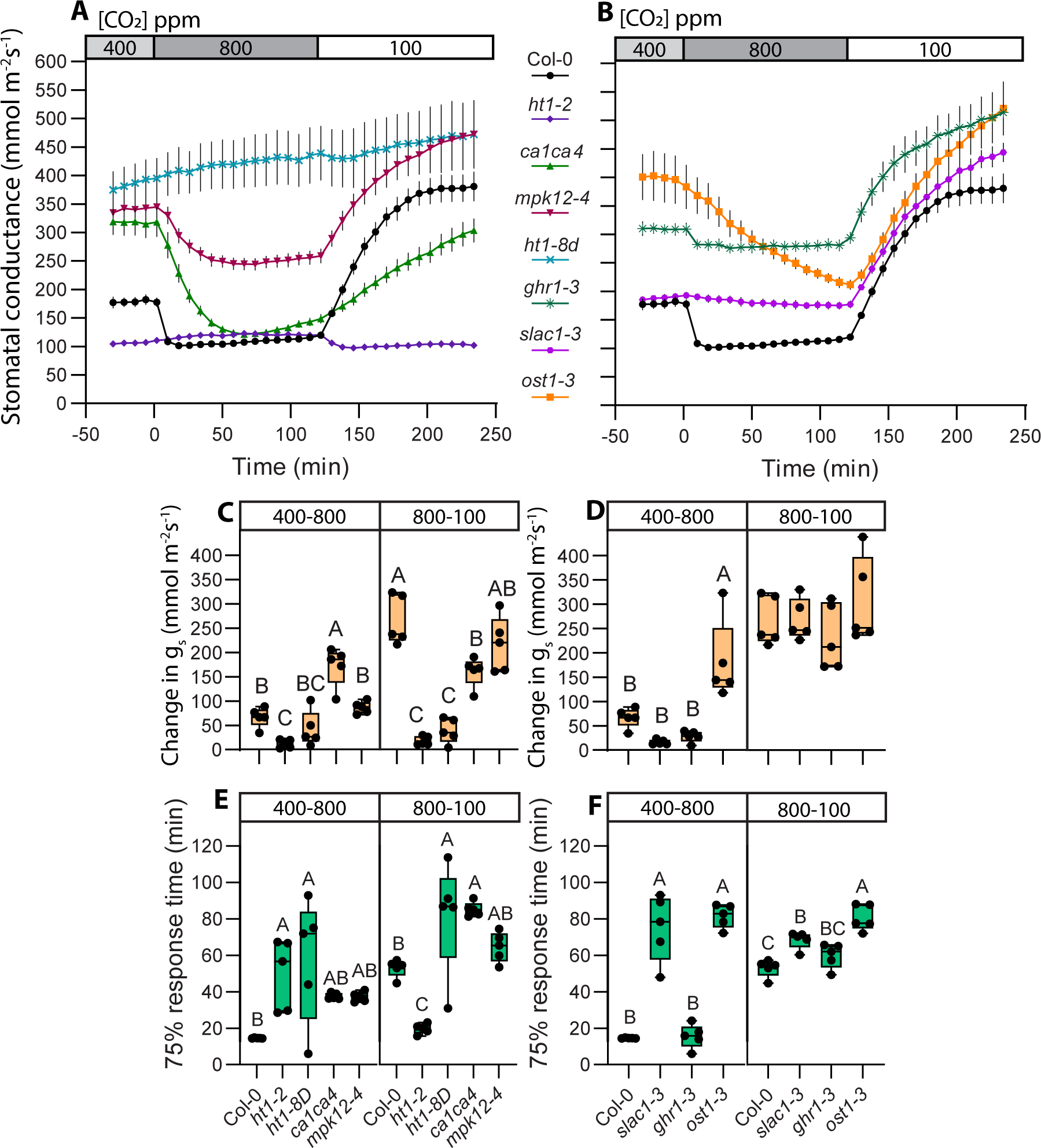
Stomatal opening in 800 to 100 ppm CO_2_ transition masks different responses present in sub-ambient and above ambient CO_2_ concentrations. **(A)** and **(B)** Stomatal response to CO_2_ concentration changes from 400 to 800 ppm and 800 to 100 ppm, mean stomatal conductance ± SEM is shown. **(C,D)** Boxplot of stomatal conductance (g_s_) change (mmol m^-2^ s^-1^) in response to CO_2_ concentration changes from 400 to 800 ppm and 800 to 100 ppm, respectively. **(E,F)** Boxplot of 75% response time (minutes) of stomatal response to CO_2_ concentration changes from 400 to 800 ppm and 800 to 100 ppm, respectively. **(C-F)** Boxes represent 25-75 % quartiles and median as the horizontal lines, whiskers indicate the smallest and largest values, points show individual plant values. Statistically significantly different groups are marked with different letters (One-way ANOVA with Tukey *post hoc* test). **(A-F)** Sample size was 5 for all plant lines. VPD during experiments was 0.9 kPa in **(A-F)**. Start of first treatment was between 11:30 to 12:30. Col-0 data is same as used in figure 1, experiments with Col-0 and mutant lines were done together.

We combined information from previously analyzed CO_2_ transitions in the mutants used in our study with our results (Table 1). Our findings mostly confirm previous results, where available, with the exception of some differences in response amplitude in the *slac1-3*, *ost1-3,* and *ca1ca4* mutants that can be explained by longer treatment duration in this work that allowed the slower stomatal responses of these mutants to reach amplitudes similar to wild-type plants. We also found faster stomatal response to sub-ambient to above-ambient CO_2_ transition in the *ht1-2*, potentially explained by different parameters used to describe response speed in different studies. Our experiments add new information on the ambient to sub-ambient, above-ambient to ambient, sub-ambient to above-ambient, above-ambient to sub-ambient and sub-ambient to ambient CO_2_ transitions that have not been systematically addressed before in all the mutants analyzed here.

**Table 1.**
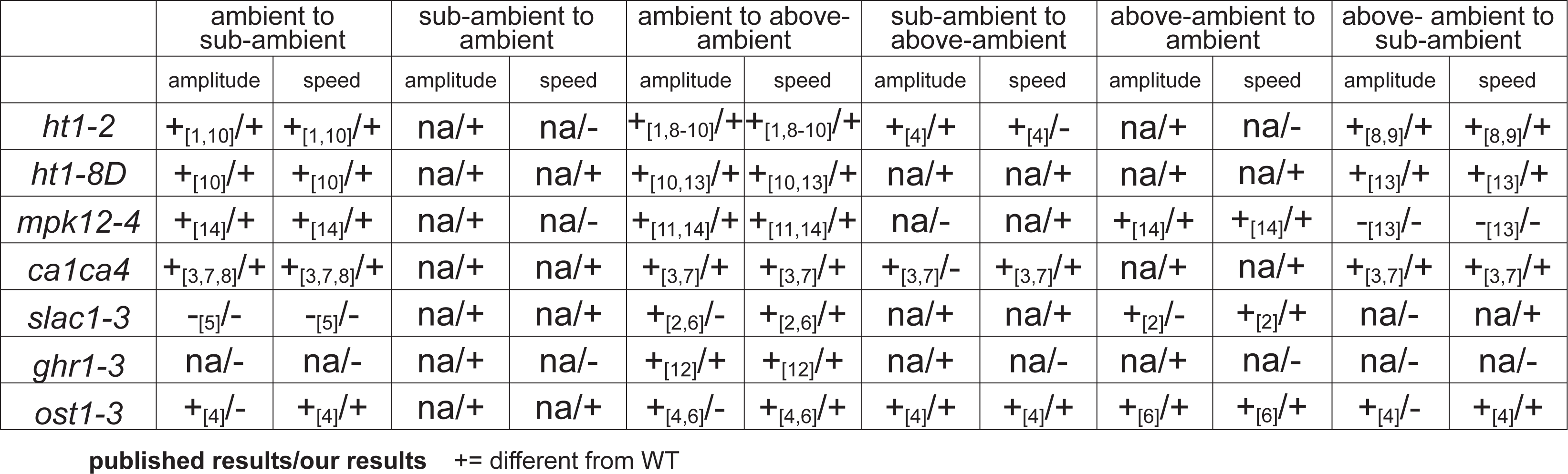
Stomatal CO2 responses for the studied mutants from the previously published studies (Hashimoto et al., 2006 [1]; Vahisalu et al., 2008 [2]; Hu et al., 2010 [3]; Xue et al., 2011 [4]; Laanemets et al., 2013 [5]; Merilo et al., 2013 [6]; Hu et al., 2015 [7]; Matrosova et al., 2015 [8]; Hashimoto-Sugimoto et al., 2016 [9]; Hõrak et al., 2016 [10]; Jakobson et al., 2016 [11]; Sierla et al., 2018 [12]; Takahashi et al., 2022 [13]; Yeh et al., 2023 [14]) and from this study.

## Discussion

Here we show that stomatal closure in response to an increase in CO_2_ concentration, which occurs both at sub-ambient and above-ambient CO_2_ concentration ranges, is regulated by both the CO_2_-specific and SLAC1-related pathways under all CO_2_ ranges, but these pathways have a different degree of importance under different CO_2_ concentration ranges. Stomatal closure in response to the 400-800 CO_2_ concentration range is faster (Fig. 1B), while the 100-400 response has greater amplitude (Fig. 1A, 2C). These differences may be explained by higher guard cell volume and larger stomatal apertures under lower CO_2_ levels, leading to slower responses with larger amplitudes, similar to slower responses of larger stomata (Drake et al., 2013; Kübarsepp et al., 2020). However, the relative contribution of the components involved in stomatal CO_2_-signalling is also different in the 400-800 and 100 to 400 stomatal closure responses (Figures 2-3). Increasing CO_2_ abruptly from 100 to 800 masked the presence of two processes with different kinetics (Figures 2, 3 and 4), indicating the necessity to analyze stomatal closure responses to elevated CO_2_ levels separately in the ambient to above-ambient and sub-ambient to ambient CO_2_ concentration ranges.

The HT1 kinase is required for plant stomatal CO_2_ signaling; plants with impaired HT1 function are nearly insensitive to all CO_2_ concentration changes (Hashimoto et al., 2006; Hashimoto-Sugimoto et al., 2016; Hõrak et al., 2016). HT1 together with MPK12 or MPK4 forms a primary CO_2_ sensing complex, where CO_2_/bicarbonate triggers interaction of MPKs with HT1 and this leads to inhibition of the HT1 kinase activity (Takahashi *et al*., 2022). In our experiments, *mpk12-4* plants had disrupted stomatal response amplitude to the 100-400 CO_2_ transition, while the 400-800 amplitude was unaffected (Fig. 2A,C). In Tõldsepp *et al,* (2018) *mpk12 mpk4GC* double-mutants, where MPK4 expression is suppressed only in guard cells, had no 400-800 CO_2_ response, yet *mpk4GC* single-mutants responded to the 400-800 CO_2_ transition similar to wild-type plants. Takahashi *et al*. (2022) showed MPK12 mutants to have the 400-800 response similar to current study, although neither of these studies tested the 100-400 response. Thus, it seems that the MPK12/MPK4-HT1 complex largely loses functionality if HT1 is impaired (Fig. 2), same happens if both MPK12 and MPK4 are missing from guard cells (Tõldsepp et al., 2018), but losing only MPK12 preferentially affects the 100-400 CO_2_ transition (Fig. 2). Therefore, while in the 400-800 transition the lack of MPK12 is likely compensated by MPK4, MPK4 cannot effectively replace the function of MPK12 at sub-ambient CO_2_ levels. This could mean that while MPK12 and MPK4 both can form CO_2_/bicarbonate sensing complex with HT1, their affinity for CO_2_/bicarbonate may be different.

CA1 and CA4 also affected stomatal responsiveness more in the 100-400 CO_2_ range, with slow and shallow stomatal response in the *ca1ca4* mutant (Fig. 2). The strong stomatal closure in above-ambient 400-800 CO_2_ transition in this mutant (Fig. 2) might be related to increased autonomous CO_2_ conversion to HCO_3_^-^ in the elevated CO_2_ environment due to shifting of the reaction balance towards bicarbonate production under above-ambient CO_2_ levels; or by involvement of other carbonic anhydrases, as recently demonstrated (Sun et al., 2022).

SLAC1 is important for stomatal closure responses to both elevated CO_2_ transitions, but compared with wild-type, the response speed of *slac1-3* in the 400-800 transition was more affected than in the 100-400 transition (Fig. 3A,E). In *slac1-3* plants there was small in magnitude and slow stomatal closure and opening in the 400-800-400 CO_2_ transitions, yet stomatal opening in response to the 400-100 transition was strong and similar in speed to wild-type plants (Supplementary Fig. 1B,D,F), further supporting the major role of SLAC1 in stomatal responses under ambient to above-ambient CO_2_ concentration changes. In the 100-400 CO_2_ transition, stomatal responsiveness may be partly compensated by other ion channels that are functional in *slac1-3*. In addition to the S-type anion channels like SLAC1, stomatal closure is also affected by R-type anion channels, such as QUAC1 (Meyer et al., 2010; Imes et al., 2013). Jalakas *et al*. (2021a) demonstrated that the *quac1-1 slac1-3* double mutant and *quac1-1 slac1-3 slah1-3* triple-mutant had no response to the 400-800 CO_2_ transition, while individually *slac1-3* had a weak response and *quac1-1* stomatal response was similar to wild-type plants. SLAH3 is another S-type anion channel contributing to stomatal closure (Zhang et al., 2016) and could potentially compensate for the lack of SLAC1 in the 100-400 CO_2_ response, although it does not affect the 400-800 stomatal CO_2_ response (Jalakas et al., 2021a). Future studies should address the 100-400 CO_2_ response in mutants deficient in major S- and R-type anion channels to better understand their potential role in the 100-400 CO_2_ response.

Stomatal opening triggered by decreased CO_2_ levels, from 400 to 100 and from 800 to 100, had larger magnitude than the 800-400 response (Fig. 1A,C,E,G). Responses to 100 ppm final CO_2_ concentration had a consistently large amplitude and slow response rate in different experimental set-ups and were not affected by the order of CO_2_ treatments. Thus, stomatal opening in response to sub-ambient CO_2_ concentrations is a very prominent response. This is in line with Merilo *et al*., (2014), where two stimuli with an opposing effect on stomata were simultaneously applied, such as darkness and low CO_2_, or low CO_2_ and elevated VPD. In such combinations, Arabidopsis stomata always opened in response to sub-ambient CO_2_ levels, further indicating that reduction of CO_2_ is a strong and prevailing signal. The sub-ambient CO_2_-induced stomatal opening, similar to light-induced opening, likely involves H^+^ATPase activation (Inoue and Kinoshita, 2017), changes in sugar and starch metabolism (Flütsch and Santelia, 2021), and suppression of stomatal closure via e.g. inhibiting anion channel activation (Marten et al., 2007). The combined activation of all these processes may explain the slow stomatal opening rate in response to sub-ambient CO_2_ levels.

Elevated VPD triggers ABA biosynthesis (McAdam et al., 2016), that can potentially increase stomatal responsiveness to CO_2_ due to interactions of CO_2_ and ABA signaling pathways (Raschke, 1975; Merilo et al., 2013; Chater et al., 2015). In previous experiments, elevated VPD has been shown to either increase stomatal responsiveness to elevated CO_2_, potentially via enhanced ABA levels (Bunce, 1998), or decrease it, potentially due to reduced stomatal apertures under elevated VPD (Morison and Gifford, 1983; Talbott et al., 2003). Our results showed an enhanced stomatal response speed in the 400-100, 100-400 and 800-100 CO_2_ concentration transitions but not in the already faster 400-800 transition (Fig. 1A,B,E,F). Elevated VPD has been shown to accelerate stomatal opening in light in angiosperms due to reduced back-pressure of epidermal cells on guard cells (Mott et al., 1999; Pichaco et al., 2024), our results of faster stomatal opening in response to sub-ambient [CO_2_] under elevated VPD are in line with this (Fig. 1B,E). Additionally, elevated VPD also accelerated stomatal closure responses to elevated CO_2_ levels in the sub-ambient to ambient concentration ranges in wild-type Arabidopsis (Fig. 1B). Faster stomatal closure may be caused by increased ABA levels under elevated VPD conditions (McAdam and Brodribb, 2015) or may be explained by the smaller steady-state stomatal conductance caused by smaller stomatal apertures under elevated VPD that can adjust faster in response to environmental changes.

Steady-states of stomatal conductance of all plant lines were decreased by elevated VPD (Fig. 2A,B, 3A,B), confirming that VPD is an important factor for steady-state stomatal conductance (Grossiord et al., 2020; López et al., 2021). Mutants with disrupted stomatal ABA response, *ghr1-3*, *ost1-3* and *slac1-3*, also had lower steady-state stomatal conductances under elevated VPD (Fig. 3A,B). Their high VPD-induced decrease in steady-state stomatal conductance could be caused by ABA-independent active processes (e.g. ABA-independent OST1 activation, whose contribution to stomatal closure under high VPD increases in time according to Jalakas et al., (2021b)), or hydropassive stomatal closure. These data indicate that elevated ABA levels alone are not sufficient to explain the decrease of stomatal conductance under elevated VPD conditions. CO_2_-induced stomatal responses under elevated VPD were sometimes faster and mostly had a smaller amplitude (Fig. 2B,D,F and 3B,D,F). However, response to the 400-800 CO_2_ transition disappeared completely in the *ghr1-3* plants under elevated VPD conditions (Fig. 3B,D). GHR1 contributes to stomatal closure in response to both elevated VPD (Hsu et al., 2021) and CO_2_ (Hõrak et al., 2016; Sierla et al., 2018). Thus, CO_2_ and VPD responses may interact in *ghr1-3*: if the relatively small elevated VPD-induced stomatal closure already occurred in the *ghr1-3* plants subjected to elevated VPD (Sierla et al., 2018; Hsu et al., 2021), no further response to CO_2_ elevation was triggered.

Here we have focused on guard cell-specific stomatal CO_2_ signaling components and their different contribution to stomatal closure responses in the sub-ambient to ambient and ambient to above-ambient CO_2_ concentration transitions. In addition to these components, it is likely that signal mediators outside guard cells also contribute to different stomatal CO_2_ response characteristics under different CO_2_ levels. While guard cells in isolated epidermis can respond to elevated CO_2_ levels by a decrease in stomatal aperture (Webb et al., 1996; Chater et al., 2015), signals from mesophyll are needed for strong CO_2_-induced stomatal closure (Mott et al., 2008; Fujita et al., 2013). Mesophyll processes, e.g. photosynthesis and sugar metabolism, are known to impact on stomatal behavior (Lawson and Matthews, 2020), and their contribution to different stomatal CO_2_ response patterns under different CO_2_ concentration ranges merits further study.

Elevated temperatures caused by climate change increase evaporative demand of the atmosphere manifested as higher VPD levels, which increases transpiration, and triggers stomatal closure to avoid wilting. Together, changes in atmospheric VPD and CO_2_ levels are perhaps the greatest agricultural challenges of the future, yet how both these stimuli together affect plant stomatal behavior is poorly understood. Here we show that while elevated VPD negatively affects steady-state stomatal conductances, it has little effect on stomatal CO_2_-responsiveness. Nevertheless, in some genetic backgrounds, we found an interaction between CO_2_ and VPD treatments, indicating that the simultaneous effects of these factors on stomatal behavior merit further study. We also show that stomatal response to elevated CO_2_ has different kinetics under sub-ambient and above-ambient CO_2_ concentration ranges and its known regulators contribute to a different degree under different CO_2_ concentration transitions. Thus, to better understand stomatal responses to CO_2_ it is necessary to carefully consider CO_2_ levels and experimental set-ups.

## Acknowledgements

This work was supported by the Estonian Research Council grant PSG404 to H.H., Basic funding from the Institute of Technology, PRG719, and PRG1620 to E.M. and PRG433 to H.K.; and European Regional Development Fund via Center of Excellence in Molecular Cell Engineering.

## Author Contributions

H.H. designed the study, H.H., K.K. and E.M. performed experiments, H.H., K.K., E.M. and H.K. analyzed data, H.H., K.K. and H.K. wrote the manuscript, all authors commented, edited and approved the final manuscript.

## References

Azoulay-Shemer T, Palomares A, Bagheri A, Israelsson-Nordstrom M, Engineer CB, Bargmann BOR, Stephan AB, Schroeder JI (2015) Guard cell photosynthesis is critical for stomatal turgor production, yet does not directly mediate CO_2_- and ABA-induced stomatal closing. Plant J 83: 567–581

Brandt B, Brodsky DE, Xue S, Negi J, Iba K, Kangasjärvi J, Ghassemian M, Stephan AB, Hu H, Schroeder JI (2012) Reconstitution of abscisic acid activation of SLAC1 anion channel by CPK6 and OST1 kinases and branched ABI1 PP2C phosphatase action. Proc Natl Acad Sci 109: 10593–10598

Brodribb TJ, McAdam SAM, Jordan GJ, Feild TS (2009) Evolution of stomatal responsiveness to CO_2_ and optimization of water-use efficiency among land plants. New Phytol 183: 839–847

Chater C, Peng K, Movahedi M, Dunn JA, Walker HJ, Liang Y-K, McLachlan DH, Casson S, Isner JC, Wilson I, et al (2015) Elevated CO_2_-Induced Responses in Stomata Require ABA and ABA Signaling. Curr Biol 25: 2709–2716

Cutler SR, Rodriguez PL, Finkelstein RR, Abrams SR (2010) Abscisic Acid: Emergence of a Core Signaling Network. Annu Rev Plant Biol 61: 651–679

Flütsch S, Santelia D (2021) Mesophyll-derived sugars are positive regulators of light-driven stomatal opening. New Phytol 230: 1754–1760

Franks PJ, Britton-Harper ZJ (2016) No evidence of general CO_2_ insensitivity in ferns: one stomatal control mechanism for all land plants? New Phytol 211: 819–827

Hammer O, Harper DAT, Ryan PD (2001) PAST: Paleontological Statistics Software Package for Education and Data Analysis. Palaeontol Electron 4: 10

Hashimoto M, Negi J, Young J, Israelsson M, Schroeder JI, Iba K (2006) Arabidopsis HT1 kinase controls stomatal movements in response to CO_2_. Nat Cell Biol 8: 391–397

Hashimoto-Sugimoto M, Negi J, Monda K, Higaki T, Isogai Y, Nakano T, Hasezawa S, Iba K (2016) Dominant and recessive mutations in the Raf-like kinase *HT1* gene completely disrupt stomatal responses to CO_2_ in Arabidopsis. J Exp Bot 67: 3251–3261

Hayashi M, Sugimoto H, Takahashi H, Seki M, Shinozaki K, Sawasaki T, Kinoshita T, Inoue S (2020) Raf-like kinases CBC1 and CBC2 negatively regulate stomatal opening by negatively regulating plasma membrane H + - ATPase phosphorylation in Arabidopsis. Photochem Photobiol Sci 19: 88–98

Hiyama A, Takemiya A, Munemasa S, Okuma E, Sugiyama N, Tada Y, Murata Y, Shimazaki K (2017) Blue light and CO 2 signals converge to regulate light-induced stomatal opening. Nat Commun 8: 1–13

Hõrak H, Kollist H, Merilo E (2017) Fern Stomatal Responses to ABA and CO_2_ Depend on Species and Growth Conditions. Plant Physiol 174: 672–679

Hõrak H, Sierla M, Tõldsepp K, Wang C, Wang Y-S, Nuhkat M, Valk E, Pechter P, Merilo E, Salojärvi J, et al (2016) A Dominant Mutation in the HT1 Kinase Uncovers Roles of MAP Kinases and GHR1 in CO_2_-Induced Stomatal Closure. Plant Cell 28: 2493–2509

Hsu P-K, Takahashi Y, Merilo E, Costa A, Zhang L, Kernig K, Lee KH, Schroeder JI (2021) Raf-like kinases and receptor-like (pseudo)kinase GHR1 are required for stomatal vapor pressure difference response. Proc Natl Acad Sci 118: e2107280118

Hu H, Boisson-Dernier A, Israelsson-Nordström M, Böhmer M, Xue S, Ries A, Godoski J, Kuhn JM, Schroeder JI (2010) Carbonic anhydrases are upstream regulators of CO_2_-controlled stomatal movements in guard cells. Nat Cell Biol 12: 87–93

Hua D, Wang C, He J, Liao H, Duan Y, Zhu Z, Guo Y, Chen Z, Gong Z (2012) A Plasma Membrane Receptor Kinase, GHR1, Mediates Abscisic Acid- and Hydrogen Peroxide-Regulated Stomatal Movement in *Arabidopsis*. Plant Cell Online 24: 2546–2561

Imes D, Mumm P, Böhm J, Al-Rasheid KAS, Marten I, Geiger D, Hedrich R (2013) Open stomata 1 (OST1) kinase controls R–type anion channel QUAC1 in Arabidopsis guard cells. Plant J 74: 372–382

Inoue S, Kinoshita T (2017) Blue Light Regulation of Stomatal Opening and the Plasma Membrane H^+^-ATPase. Plant Physiol 174: 531–538

Jakobson L, Vaahtera L, Tõldsepp K, Nuhkat M, Wang C, Wang Y-S, Hõrak H, Valk E, Pechter P, Sindarovska Y, et al (2016) Natural Variation in *Arabidopsis* Cvi-0 Accession Reveals an Important Role of MPK12 in Guard Cell CO_2_ Signaling. PLOS Biol 14: e2000322

Jalakas P, Nuhkat M, Vahisalu T, Merilo E, Brosché M, Kollist H (2021a) Combined action of guard cell plasma membrane rapid- and slow-type anion channels in stomatal regulation. Plant Physiol 187: 2126–2133

Jalakas P, Takahashi Y, Waadt R, Schroeder JI, Merilo E (2021b) Molecular mechanisms of stomatal closure in response to rising vapour pressure deficit. New Phytol 232: 468–475

Katsuhara M, Hanba YT (2008) Barley plasma membrane intrinsic proteins (PIP Aquaporins) as water and CO_2_ transporters. Pflüg Arch - Eur J Physiol 456: 687–691

Kollist T, Moldau H, Rasulov B, Oja V, Rämma H, Hüve K, Jaspers P, Kangasjärvi J, Kollist H (2007) A novel device detects a rapid ozone-induced transient stomatal closure in intact Arabidopsis and its absence in abi2 mutant. Physiol Plant 129: 796–803

López J, Way DA, Sadok W (2021) Systemic effects of rising atmospheric vapor pressure deficit on plant physiology and productivity. Glob Change Biol 27: 1704–1720

Marten H, Hedrich R, Roelfsema MRG (2007) Blue light inhibits guard cell plasma membrane anion channels in a phototropin-dependent manner. Plant J 50: 29–39

McAdam SAM, Brodribb TJ (2015) The Evolution of Mechanisms Driving the Stomatal Response to Vapor Pressure Deficit. Plant Physiol 167: 833–843

Merilo E, Yarmolinsky D, Jalakas P, Parik H, Tulva I, Rasulov B, Kilk K, Kollist H (2018) Stomatal VPD Response: There Is More to the Story Than ABA. Plant Physiol 176: 851–864

Meyer S, Mumm P, Imes D, Endler A, Weder B, Al-Rasheid KAS, Geiger D, Marten I, Martinoia E, Hedrich R (2010) AtALMT12 represents an R-type anion channel required for stomatal movement in Arabidopsis guard cells. Plant J 63: 1054–1062

Sierla M, Hõrak H, Overmyer K, Waszczak C, Yarmolinsky D, Maierhofer T, Vainonen JP, Salojärvi J, Denessiouk K, Laanemets K, et al (2018) The Receptor-like Pseudokinase GHR1 Is Required for Stomatal Closure. Plant Cell 30: 2813–2837

Sun P, Isner J-C, Coupel-Ledru A, Zhang Q, Pridgeon AJ, He Y, Menguer PK, Miller AJ, Sanders D, Mcgrath SP, et al (2022) Countering elevated CO_2_ induced Fe and Zn reduction in Arabidopsis seeds. New Phytol 235: 1796– 1806

Takahashi Y, Bosmans KC, Hsu P-K, Paul K, Seitz C, Yeh C-Y, Wang Y-S, Yarmolinsky D, Sierla M, Vahisalu T, et al (2022) Stomatal CO_2_/bicarbonate sensor consists of two interacting protein kinases, Raf-like HT1 and non-kinase-activity activity requiring MPK12/MPK4. Sci Adv 8: eabq6161

Tõldsepp K, Zhang J, Takahashi Y, Sindarovska Y, Hõrak H, Ceciliato PHO, Koolmeister K, Wang Y-S, Vaahtera L, Jakobson L, et al (2018) Mitogen-activated protein kinases MPK4 and MPK12 are key components mediating CO_2_-induced stomatal movements. Plant J 96: 1018–1035

Vahisalu T, Kollist H, Wang Y-F, Nishimura N, Chan W-Y, Valerio G, Lamminmäki A, Brosché M, Moldau H, Desikan R, et al (2008) SLAC1 is required for plant guard cell S-type anion channel function in stomatal signalling. Nature 452: 487–491

Wang C, Hu H, Qin X, Zeise B, Xu D, Rappel W-J, Boron WF, Schroeder JI (2016) Reconstitution of CO_2_ Regulation of SLAC1 Anion Channel and Function of CO_2_-Permeable PIP2;1 Aquaporin as CARBONIC ANHYDRASE4 Interactor. Plant Cell 28: 568–582

Xue S, Hu H, Ries A, Merilo E, Kollist H, Schroeder JI (2011) Central functions of bicarbonate in S-type anion channel activation and OST1 protein kinase in CO_2_ signal transduction in guard cell. EMBO J 30: 1645–1658

Yeh C-Y, Wang Y-S, Takahashi Y, Kuusk K, Paul K, Arjus T, Yadlos O, Schroeder JI, Ilves I, Garcia-Sosa AT, et al (2023) MPK12 in stomatal CO_2_ signaling: function beyond its kinase activity. New Phytol 239: 146–158

Yoshida R, Hobo T, Ichimura K, Mizoguchi T, Takahashi F, Aronso J, Ecker JR, Shinozaki K (2002) ABA-Activated SnRK2 Protein Kinase is Required for Dehydration Stress Signaling in *Arabidopsis*. Plant Cell Physiol 43: 1473–1483

Zhang A, Ren H-M, Tan Y-Q, Qi G-N, Yao F-Y, Wu G-L, Yang L-W, Hussain J, Sun S-J, Wang Y-F (2016) S-Type Anion Channels SLAC1 and SLAH3 Function as Essential Negative Regulators of Inward K^+^ Channels and Stomatal Opening in Arabidopsis. Plant Cell 28: 949–965

