## Supplementary Figure 1 for "Stomatal CO_2_-responses at sub- and above-ambient CO_2_ levels employ different pathways in Arabidopsis"

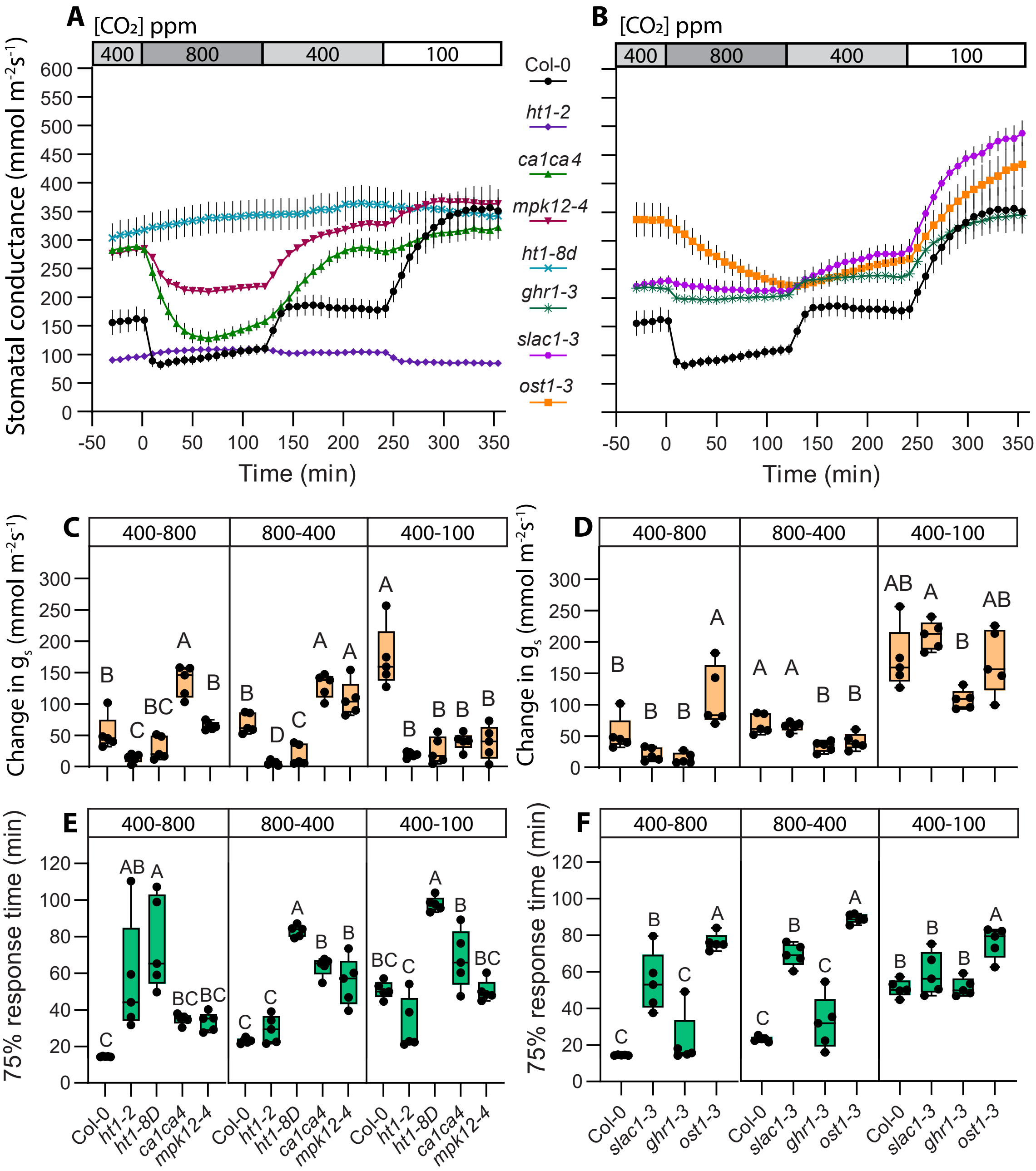


**Supplementary figure 1**

**(A)** and **(B)** Stomatal response to $CO_2$ concentration changes from 400 to 800 ppm, 800 to 400 ppm and 400 to 100 ppm, mean stomatal conductance ± SEM is shown. **(C,D)** Boxplot of stomatal conductance ($g_s$) change (mmol $m^{-2}$ $s^{-1}$) in response to $CO_2$ concentration changes from 400 to 800 ppm, 800 to 400 ppm and 400 to 100 ppm, respectively. **(E,F)** Boxplot of 75% response time (minutes) of stomatal response to $CO_2$ concentration changes from 400 to 800 ppm, 800 to 400 ppm and 400 to 100 ppm, respectively. **(C-F)** Boxes represent 25-75 % quartiles and median as the horizontal lines, whiskers indicate the smallest and largest values, points show individual plant values. Statistically significantly different groups are marked with different letters (One-way ANOVA with Tukey *post hoc* test). **(A-F)** Sample size was 5 for all plant lines. VPD during experiments was 0.9 kPa in **(A-F)**. Start of first treatment was between 11:30 to 12:30. Col-0 data is same as used in figure 1, experiments with Col-0 and mutant lines were done together.
